# Strong habitat-associated differentiation of microbial communities across a fine-scale land-water interface

**DOI:** 10.64898/2026.10.22.760677

**Authors:** Chloé Haberkorn, Christian I. Oporto, Javier Pinto, Rike Stelkens

## Abstract

Microbial communities can remain strongly differentiated across environmental boundaries despite their potential for widespread dispersal. However, the extent to which habitat differences structure microbial communities at fine spatial scales remains poorly understood. Here, we investigated bacterial and fungal communities across a 350-m land-water interface in Lake Vänern, Sweden, including mainland soil, two small terrestrial islands and lake water. Using 16S and ITS amplicon sequencing, we compared microbial diversity and community composition across habitats and with increasing distance from the shoreline. Both bacterial and fungal communities showed pronounced habitat-associated differentiation, with the strongest contrasts between terrestrial and aquatic samples. Despite being surrounded by lake water, island communities were compositionally more similar to mainland soil communities, suggesting that geographic proximity alone does not necessarily lead to microbial community homogenization. Fungal, but not bacterial community composition varied significantly with shoreline position within both terrestrial and aquatic habitats, indicating contrasting spatial patterns between the two groups. Fungal genera also showed greater taxonomic overlap between terrestrial and aquatic habitats than bacterial genera, although overall community composition remained distinct. Taxonomically inferred functional profiles and ecological guilds further reflected habitat-associated differences. Together, our findings reveal strong microbial community differentiation across a fine-scale land-water interface and highlight how habitat identity and spatial context are associated with distinct patterns of bacterial and fungal community assembly.

## Introduction

Microorganisms are a major component of biodiversity and may account for over 99% of all species (Locey and Lennon, 2016). They encompass diverse groups, including archaea, protists, bacteria and fungi, the latter two representing most of the global living biomass within microbes (Crowther *et al*., 2024). Microorganisms are found across all environments, where they mediate essential processes such as nutrient cycling and decomposition, and are part of biogeochemical cycles (Cotner and Biddanda, 2002; Van Der Heijden, Bardgett and Van Straalen, 2008).

Despite their ubiquity, microorganisms are not uniformly distributed across environments and understanding how microbial communities are organised in space has become a central question in microbial ecology (Nemergut Diana R. *et al*., 2013). Microorganisms are generally considered to have limited active dispersal, but high passive dispersal potential, partly due to their small size, which facilitates transport across environments by wind, water and other vectors (Finlay, 2002; Wilkinson *et al*., 2012). This has led to the paradigm that “*everything is everywhere: but the environment selects*”: microbial organisms disperse widely, and environmental conditions then determine where they can establish (Becking, 1934). However, subsequent studies have refined this view, by demonstrating that microbial communities can exhibit clear biogeographic structure, with microbial community composition shaped by both environmental conditions and spatial processes such as dispersal limitation (Martiny *et al*., 2006; O’Malley, 2008). Dispersal and environmental filtering can act in opposing ways. While dispersal increases the exchange of microorganisms and thereby favours community homogenization (Fodelianakis *et al*., 2019), differences in environmental conditions select for different subsets of the local species pool, strengthening spatial boundaries between communities (Evans, Martiny and Allison, 2017). The relative importance of these processes therefore depends on the geographical distance and environmental differentiation between the habitats, but also the dispersal capacities of the organisms themselves (Evans, Martiny and Allison, 2017).

Land-lake interfaces provide a particularly useful setting in which to examine this balance (Richards *et al*., 2024). Terrestrial and aquatic habitats can be highly connected, with microbial exchange occurring through runoff and atmospheric transport (Singh *et al*., 2026). Such connectivity can facilitate the exchange of microorganisms across habitat boundaries, but contrasting environmental conditions may limit establishment and persistence following dispersal, resulting in habitat-specific community composition (Monard *et al*., 2016). Understanding how sharply microbial communities remain differentiated across such interfaces can therefore provide insight into the interplay between dispersal and environmental selection at fine spatial scales.

This question is especially interesting for small freshwater islands where terrestrial habitats are surrounded by, and in close proximity to, an aquatic environment. From a biogeographic perspective, such islands could act as ecological bridges linking terrestrial and aquatic habitats, potentially hosting microbial communities composed of taxa from both environments. Alternatively, if environmental selection is sufficiently strong, islands may maintain distinct microbial assemblages despite their close proximity to the surrounding water. Previous studies of microbial island biogeography have shown that microbial communities can exhibit spatial structure across islands, with bacteria and fungi showing similar diversity patterns but being shaped by different ecological drivers (Li *et al*., 2020). However, the spatial scale at which habitat differentiation overcomes connectivity remains incompletely resolved.

Here, we investigated bacterial and fungal communities across a 350-m terrestrial-aquatic transect including mainland soil, two small islands and lake water. This spatial configuration allowed us to examine microbial community structure at a scale where geographic separation is limited but habitat boundaries remain pronounced, providing an opportunity to assess the relative influence of dispersal and environmental filtering across fine spatial gradients. Using amplicon sequencing and genus-level community analyses, we characterized the microbial diversity both across and within habitats, using transects extending progressively away from the shoreline, into both terrestrial and aquatic environments. We further examined the extent of community overlap among mainland, island and aquatic communities to assess whether islands support transitional microbial communities connecting terrestrial and aquatic microbial assemblages, or instead maintain distinct microbial communities shaped by their local environment. If dispersal across this spatially connected landscape promotes community homogenization, we would expect substantial overlap among neighbouring habitats. Conversely, strong environmental filtering would maintain differences in community composition despite their close geographic proximity.

## Material and Methods

### Sample collection

We collected water and soil samples from riparian and littoral zones of Lake Vänern in Värmland County, Sweden. Lake Vänern is Sweden’s largest lake with a surface area of 5K650 km². Sampling was conducted around *Getingbergets badplats* (Getingberget bathing place), near Karlstad, on Saturday, July 6th 2024, at 12°C, 80% humidity, and 5m/s SSW wind. The study area expanded across a 350-m transect comprising mainland, lake and two small island habitats. Nine sampling sites were established in each of the mainland (Land) and aquatic (Lake) habitats, along transects extending away from the land-water boundary (**Figure 1**). Within each habitat, sampling sites were distributed across three position classes: closest to the shoreline (Shore), at an intermediate distance (Middle), and furthest from the shoreline (Furthest). Three spatially separated sites were sampled within each position class, arranged from northeast to southwest (sites 1-3), resulting in nine sites per habitat. Soil was additionally sampled on two small islands within the aquatic sampling area (Island 1 and Island 2), resulting in a total of 11 terrestrial sampling sites (Figure 1).

**Figure 1.**
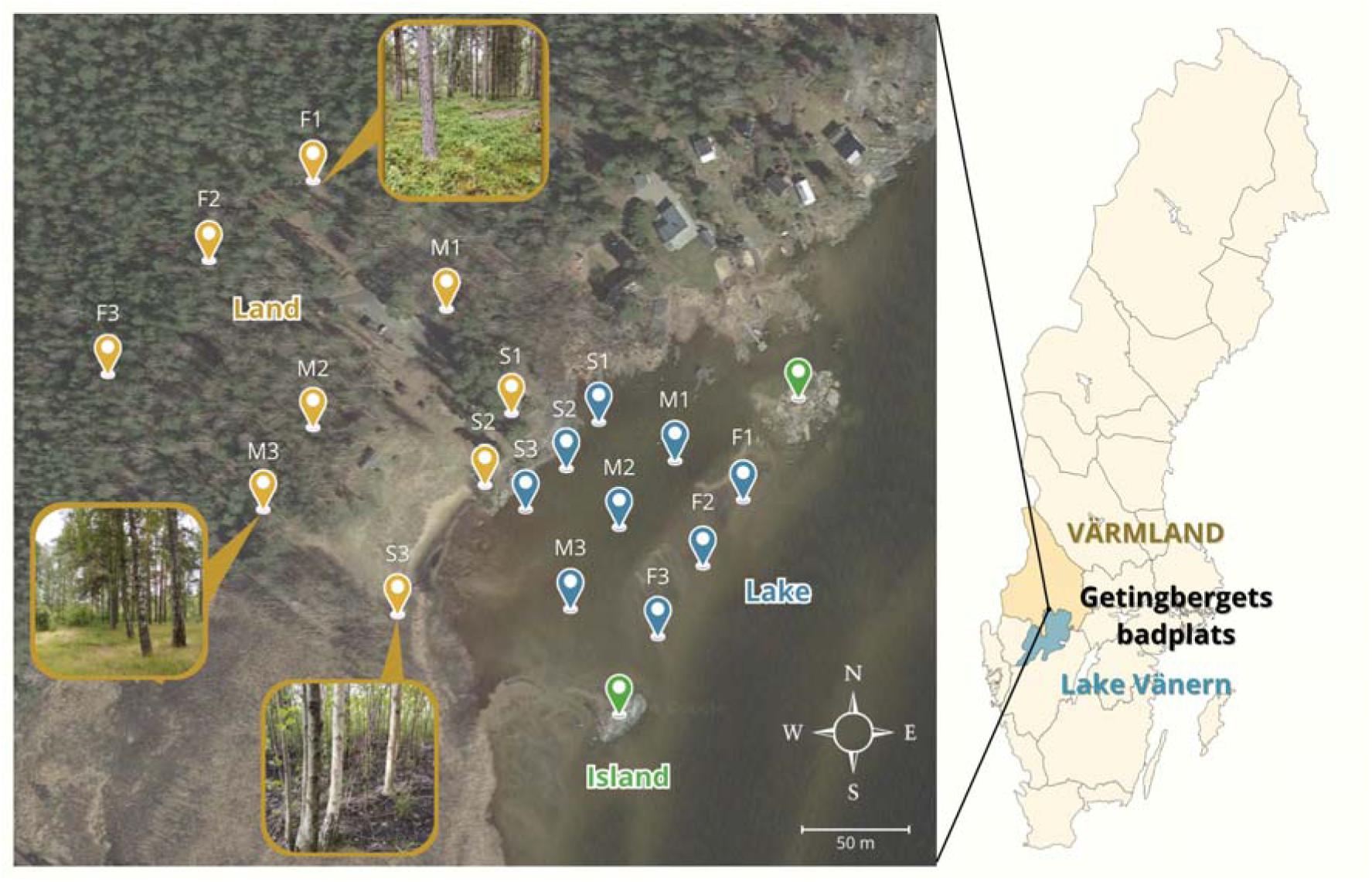
Study area and sampling design. The right panel shows the location of *Getingbergets badplats* (Getingberget bathing place) on the northern shore of Lake Vänern, Sweden. The left panel shows mainland and island (soil) sampling sites and lake (water) sampling sites. Mainland and lake sites were arranged along opposing transects extending away from the land-water boundary, with three sampling positions at increasing distances from the shoreline (S = Shore, M = Middle, and F = Furthest) and three spatially separated sites within each position (1-3). Soil was also sampled on two small islands (Island 1 and Island 2). Photographs illustrate representative vegetation at the Shore, Middle and Furthest terrestrial positions. Scale bar = 50 m.

At each terrestrial sampling site, two soil samples were extracted from a depth of 5 cm using a shovel that was thoroughly decontaminated with ethanol between samples. Soil samples were stored in a zip-lock polyethylene bag. At each aquatic sampling site, water was collected at two depths (10 and 30 cm below the water surface) using a PVC tube fitted with vinyl tubing and connected to a syringe and transferred directly into sterile 50-mL Falcon tubes (following Bhatnagar *et al*., 2020). Total water depth ranged from ca. 50 cm depth at Shore and Middle sites to 1 m at Furthest sites. The sampling equipment was thoroughly cleaned with ethanol between samples.

Furthest terrestrial sites were located in coniferous forest dominated by *Pinus*, with a ground layer composed primarily of mosses and *Vaccinium*. In contrast, Middle and Shore sites were situated in predominantly deciduous forest, dominated by *Betula*, with *Poaceae* common in the ground vegetation (see pictures in **Figure 1**). All samples were stored at 4°C until environmental DNA (eDNA) extraction.

### Environmental DNA extraction and marker amplification

Prior to DNA extraction, water samples were enriched in liquid medium consisting of yeast nitrogen base (YNB) supplemented with 1% (w/v) raffinose. For each sample, 25 mL of water was mixed with 25 mL enrichment medium in a 50 mL Falcon tube, and incubated for three days at room temperature. Following enrichment, 250 μL of the resulting culture was used for DNA extraction. For soil samples, 250 mg of soil was weighted from each sample bag for DNA extraction.

Extractions were performed using the DNeasy PowerSoil Pro Kit (QIAGEN), following the manufacturer’s protocol, with a final elution volume of 50 μL for soil samples and 100 μL for water samples. Extracted DNA from each sample was subjected to separate PCR amplifications targeting fungal and bacterial communities. For fungi, the ITS1 region was amplified using ITS1 and ITS2 primers (White *et al*., 1990; Alsammar *et al*., 2019). Bacterial 16S rRNA genes were amplified using primers targeting the V3-V4 region (Herlemann *et al*., 2011) (primer sequences are provided in **Supp. Table S1**). Amplifications were performed using the Takara Taq polymerase kit. PCR products were quantified using Qubit dsDNA HS Assay Kits and normalised to 2.5 ng/μL with Milli-Q water to a final volume of 30 μL per sample.

### Sequencing

Normalised PCR1 amplicons were purified using MagSI-DNA NGS PREP Plus magnetic beads at the National Genomics Infrastructure (NGI, Solna, Sweden). Purified amplicons were subjected to a second PCR (PCR2) for indexing with Adaptamera indices, followed by a magnetic bead purification. The final libraries were quality controlled before paired-end sequencing (2×300 bp) on the Illumina NextSeq 2000 platform. Raw reads were deposited in NCBI’s Sequence Read Archive (SRA) under BioProject accession PRJNA1251841.

### Read processing

Raw sequencing reads were processed using the nf-core/ampliseq pipeline (Straub *et al*., 2020) with default parameters and the ‘illumina_pe_its’ option for ITS amplicons. Amplicon Sequence Variants (ASVs), defined as unique amplicon sequences resolved from the sequencing data, were inferred using DADA2 (Callahan *et al*., 2016). Taxonomic assignment was performed separately for the two markers datasets. For 16S rRNA gene amplicons, ASVs were taxonomically classified using the assignTaxonomy function against the SILVA reference database, release 138.2 (Quast *et al*., 2013). The minBoot parameter was set to 70, applying a more conservative threshold than the default value of 50, following Geraldi *et al*. (2020). For ITS amplicons, taxonomic identities were assigned using BLASTn v2.16.0 with the megablast algorithm against the NCBI RefSeq Targeted Loci ITS database (ITS_eukaryote_sequences; NCBI Resource Coordinators, 2018; Camacho *et al*., 2009). The best-supported database match was retained for each ASV, and assignments with less than 97% sequence identity to the corresponding reference sequence were excluded from subsequent analyses (Nilsson *et al*., 2019).

For both marker sets, ASVs lacking genus-level taxonomic assignments were excluded from the analysis, as were ASVs represented by fewer than 10 reads across all samples or detected in only a single sample. Following filtering, 4,959 ITS ASVs and 9,699 16S ASVs were retained for downstream analyses (**Supp. Table S2 and S3**).

### Functional and ecological guild assignment

Fungal ecological guilds were assigned FUNGuild through the package R/FUNGuildR v0.4.1 (Nguyen *et al*., 2016). Assignments with a FUNGuild confidence ranking of “Probable” or “Highly Probable” were retained for downstream analyses, whereas assignments classified as “Possible” or lacking a confidence ranking were considered unassigned. Bacterial functional categories were inferred using FAPROTAX v1.2.12 through the package R/microeco v2.3.0 (Louca, Parfrey and Doebeli, 2016; Liu *et al*., 2021). ASVs assigned to multiple functional categories were included in each corresponding category.

For each sampling site (detailed in **Suppl. Table S4**), replicate read counts were summed, prior to the computation of relative abundances of fungal ecological guilds and bacterial functional categories. Categories reaching at least 1% relative abundance in at least one sampling site were retained for visualisation. The resulting relative-abundance matrices were log10-transformed after adding a pseudocount of 10^-4^ to allow the transformation of zero values. Heatmaps were generated using the package R/pheatmap v1.0.13 (Kolde and Kolde, 2015).

### Statistical analysis

Genus-level read counts were used to compute alpha diversity indices: genus richness (the number of different genus), Shannon diversity (accounting for both genus richness and the proportional abundance of each genus within a sample; Shannon & Weaver, 1963), and Simpson diversity indices (accounting for both richness and evenness, i.e., how evenly distributed genus abundances are distributed within a community; Wilsey and Potvin, 2000), using the package R/vegan v2.6-8 (Oksanen *et al*., 2020). Because the two samples collected at each site represented independent biological replicates, alpha diversity indices were calculated separately for each replicate and then averaged across replicates within each sampling site for statistical analyses.

Differences in community composition among samples were visualized using principal coordinates analysis (PCoA) based on Bray-Curtis dissimilarities calculated from genus-level abundance data, using the pcoa function in R/ape v.5.8-1.

Differences in community composition among habitats, sampling positions relative to the shoreline and sampling sites were tested using permutational multivariate analysis of variance (PERMANOVA). Bray-Curtis dissimilarity matrices were calculated from genus-level abundance data using the vegdist function from the package R/vegan (method “bray”). The two biological replicates from each sampling site were averaged after conversion to relative abundance, resulting in one community profile per site. Homogeneity of multivariate dispersion among habitats was first assessed using PERMDISP implemented with the betadisper function (R/vegan). Differences in dispersion were evaluated using the associated permutation test through the function permutest, based on 999 permutations. PERMANOVAs were then performed using the adonis2 function (R/vegan) with 999 permutations. Analyses tested (i) differences among Land, Island, and Lake habitats; (ii) differences in position relative to the shoreline (Shore, Middle, Furthest) within Land and Lake; (iii) the southwest-to-northeast spatial site axis; and (iv) the interaction between habitat (Land vs Lake) and shoreline position. For significant overall PERMANOVA tests, pairwise PERMANOVA comparisons were performed, with p-values adjusted for multiple testing using the Benjamini-Hochberg correction.

To examine associations between bacterial and fungal genera, the two biological replicates from each sampling site were summed before calculating relative abundances. Genera exceeding 1% relative abundance in at least two sampling sites were retained for network analysis. Spearman rank correlations were calculated between all pairs of retained bacterial and fungal genera using the rcorr function in the package R/Hmisc v5.2-6.

Associations were considered significant when the absolute correlation coefficient (ρ) was ≥ 0.8 and the Benjamini-Hochberg-adjusted p-value was < 0.05. The resulting network was exported in GraphML format and visualised using GephiLite.

## Results

### Alpha diversity is lower in water than in soil

Across all samples, we detected 645 bacterial and 608 fungal genera. We first analysed the within-sample diversity across sampling environments.

For bacterial genus, richness, Shannon and Simpson diversity differed significantly among Land, Island and Lake samples (**Figure 2**, Kruskal-Wallis tests; Richness: X² = 15.57, p = 4.16e-04; Shannon: X² = 14.30, p = 7.85e-04; Simpson: X² = 14.16, p = 8.42e-04). All three diversity metrics were significantly lower in Lake samples than in Land samples, while only genus richness was significantly lower in Lake samples than in Island samples (pairwise Wilcoxon rank sum test with Benjamini-Hochberg correction: p_adj_ < 0.05). Island samples also had significantly higher genus richness than Land samples (p_adj_ = 0.045), whereas Shannon and Simpson diversity did not differ significantly between these two habitats.

**Figure 2.**
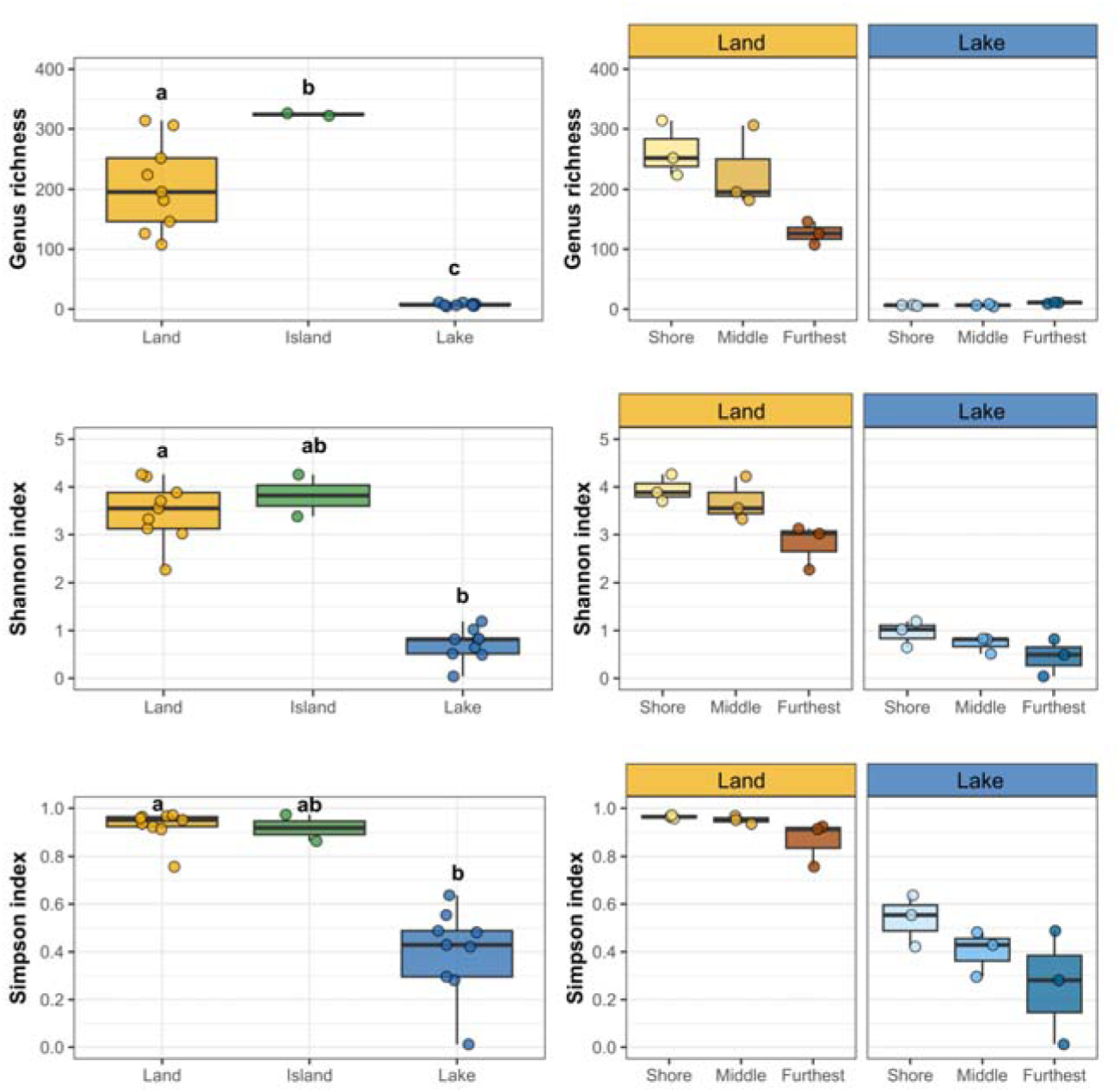
Bacterial alpha diversity across habitats and distance from the shoreline. Genus richness, Shannon and Simpson diversity are shown. Left panels compare mainland soil (Land), island soil (Island) and lake water (Lake); right panels compare Shore, Middle and Furthest positions separately within Lake and Land habitats. Points represent sampling sites, and boxplots show medians and interquartile ranges. Within each panel, groups not sharing a letter differ significantly in Benjamini-Hochberg-adjusted pairwise Wilcoxon tests following a significant Kruskal-Wallis test.

Within terrestrial samples, bacterial diversity declined with increasing distance from the shoreline, although sites differ only marginally significantly from each other (Kruskal-Wallis tests; Richness, Shannon and Simpson: X² = 5.96, p = 0.051). Within lake samples, although the same trend seemed to be observed for Simpson diversity, none of the three diversity metrics differed significantly with distance from shore (all p > 0.066).

Fungal communities showed a similar overall pattern to bacterial communities. Genus richness, Shannon diversity and Simpson diversity differed significantly among Land, Island and Lake samples (**Figure 3**, Kruskal-Wallis test; Richness: X² = 14.21, p = 8.20e-04; Shannon: X² = 12.84, p = 1.63e-03; Simpson: X² = 11.70, p = 2.88e-03). All three diversity metrics were significantly higher in Land samples than in Lake samples (pairwise Wilcoxon rank sum test with Benjamini-Hochberg correction: all p_adj_ < 8.60e-03), while Island samples showed higher diversity than Lake samples, although this difference did not reach the significance threshold (all p_adj_ = 0.055). Unlike bacterial communities, Land and Island samples did not differ significantly in genus richness (p = 0.73).

**Figure 3.**
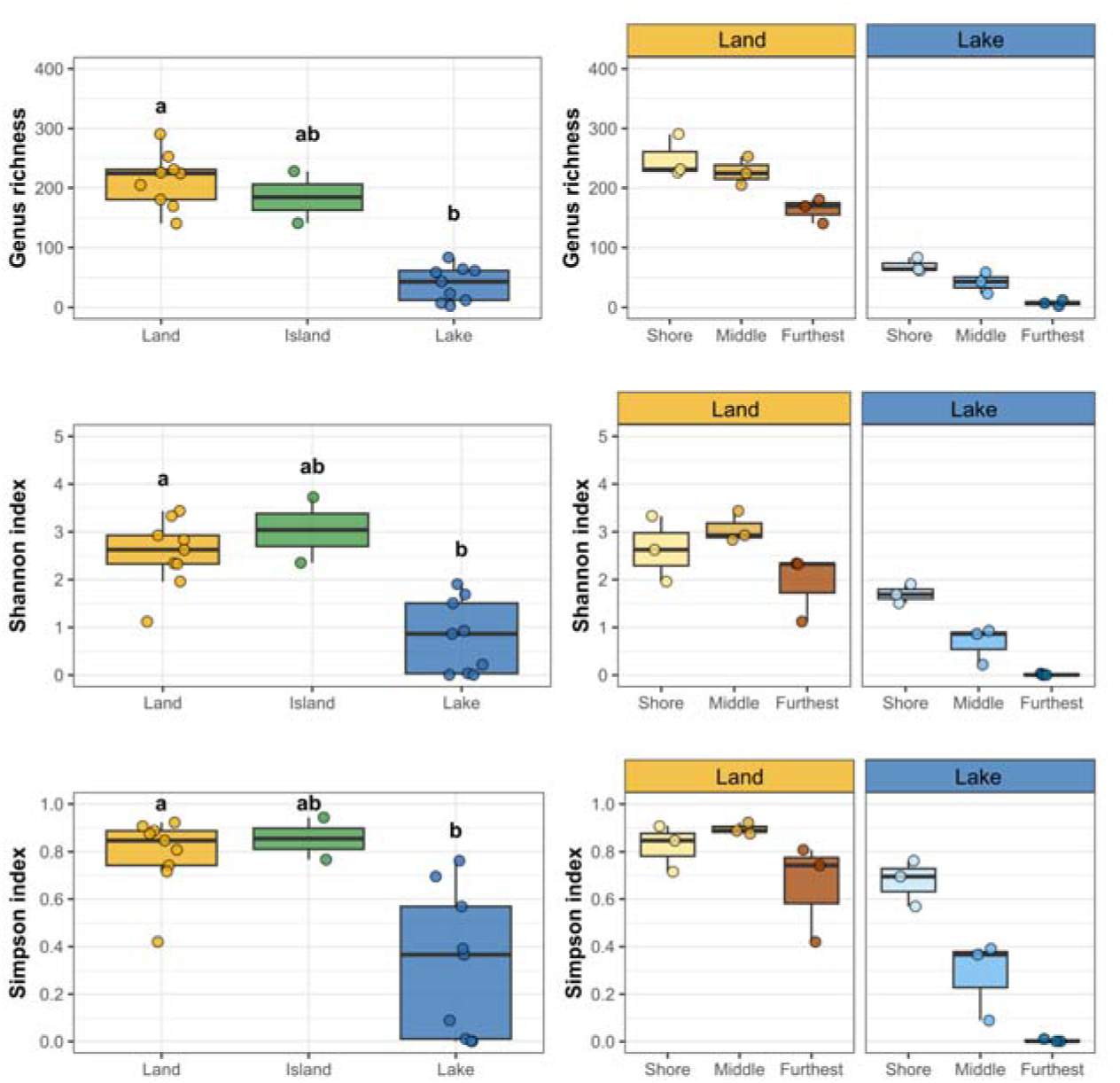
Fungal alpha diversity across habitats and distance from the shoreline. Genus richness, Shannon and Simpson diversity are shown. Left panels compare mainland soil (Land), island soil (Island) and lake water (Lake); right panels compare Shore, Middle and Furthest positions separately within Lake and Land habitats. Points represent sampling sites. Boxplots show medians and interquartile ranges. Within each panel, groups not sharing a letter differ significantly in Benjamini-Hochberg-adjusted pairwise Wilcoxon tests following a significant Kruskal-Wallis test.

Within terrestrial habitats, genus richness decreased with increasing distance from the shoreline, whereas other alpha diversity indexes were on average higher in Middle sites. However, none of the pairwise comparisons between sampling sites were statistically significant (pairwise Wilcoxon rank sum tests with Benjamini-Hochberg correction, all p_adj_ > 0.15). Within the Lake, the decline in fungal alpha diversity with increasing distance from the shoreline was more consistent across diversity metrics. However, as in terrestrial habitats, none of the pairwise comparisons between sampling sites were statistically significant (all p_adj_ > 0.10).

Taken together, these analyses show that the strongest alpha diversity contrast occurred between terrestrial and aquatic habitats with substantially higher genus richness and diversity in soil than in water samples, and a putative higher diversity in the samples at the interface between the terrestrial and aquatic habitats (Shore locations). However, alpha diversity metrics do not reveal whether communities are composed of the same genera or share similar relative-abundance structures. We therefore next tested for differences in community composition among samples using Bray-Curtis dissimilarities.

### Community composition is structured primarily across the soil-water boundary

Bacterial community composition differed strongly among Land, Island, and Lake water samples (PERMANOVA based on Bray-Curtis distances, R² = 0.61, F = 13.5, p = 0.001), i.e., habitat explained over 60% of the variation in community composition. Pairwise comparisons showed that this difference was primarily driven by the strong differentiation between Land and Lake communities (Benjamini-Hochberg correction, R² = 0.62, F = 26.3, p_adj_ = 0.003) and between Island and Lake communities (R² = 0.45, F = 7.43, p_adj_ = 0.039), whereas Land and Island communities did not differ significantly (R² = 0.20, F = 2.31, p_adj_ = 0.055). The PCoA based on Bray-Curtis dissimilarities also showed a clear separation between Land and Lake samples, while the two Island sites differed in their positions: one was separated from the other samples along PC1, whereas the other clustered with the Land samples (**Supp. Figure S1**). Multivariate dispersion did not differ significantly among habitats (F = 0.99, p = 0.395), indicating no detectable difference in within-habitat variability among Land, Island, and Lake communities.

Community composition did not vary significantly among sampling sites along the southwest-to-northeast axis (R² = 0.025, F = 0.47, p = 0.774), nor with distance from the shoreline in either terrestrial or aquatic habitats (Shore, Middle, Furthest; Land: R² = 0.32, F = 1.38, p = 0.186; Lake: R² = 0.11, F = 0.39, p = 0.845). The interaction between habitat and shoreline-position was also not significant (R² = 0.042, F = 0.85, p = 0.453).

Fungal communities showed a similar overall pattern. Community composition differed significantly among Land, Island, and Lake habitats (PERMANOVA based on Bray-Curtis distances, R² = 0.44, F = 6.68, p = 0.001). Pairwise comparisons indicated that this difference was primarily driven by the contrast between Land and Lake communities (R² = 0.42, F = 11.5, p_adj_ = 0.003) and Island and Lake communities (R² = 0.37, F = 5.36, p_adj_ = 0.039) whereas differences between Land and Island communities (R² = 0.12, F = 1.18, p_adj_ = 0.275) were not significant. Multivariate dispersion did not differ significantly among habitats (F = 0.91, p = 0.412).

Community composition did not vary significantly among sampling sites along the southwest-to-northeast axis (R² = 0.031, F = 0.57, p = 0.796). However, in contrast to bacterial communities, fungal community composition varied significantly with shoreline position within both terrestrial (R² = 0.36, F = 1.67, p = 0.046) and Lake habitats (R² = 0.59, F = 4.24, p = 0.007). Further pairwise comparisons between shoreline positions did not identify any significant differences (Benjamini-Hochberg correction, all p_adj_ > 0.15), although the Furthest–Shore comparison within Lake habitats showed a relatively large effect size (R² = 0.65, F = 7.31, p_adj_ = 0.15). The interaction between habitat and shoreline-position interaction was also significant (R² = 0.13, F = 2.42, p = 0.032), indicating that the association between shoreline position and fungal community composition differed between Land and Lake habitats. We therefore further investigated the extent to which genera were shared among Land, Island and Lake communities.

### Islands communities predominantly overlap with terrestrial communities

Genus-level comparisons revealed similar habitat-association patterns for bacterial and fungal communities (Figure 4). In both datasets, terrestrial and aquatic communities were strongly differentiated, while Island communities overlapped predominantly with terrestrial communities. For bacteria, 346 genera (53.6% of all detected bacterial genera) were shared between Land and Island samples, but absent from the Lake, while 150 genera (23.3%) were unique to Land and 111 (17.2%) were unique to Island samples (**Figure 4A**). In contrast, only 38 genera were detected in Lake samples, of which 26 occurred across all three habitats. Only two genera were shared exclusively between Land and Lake and eight between Island and Lake, while two genera were unique to the Lake.

**Figure 4.**
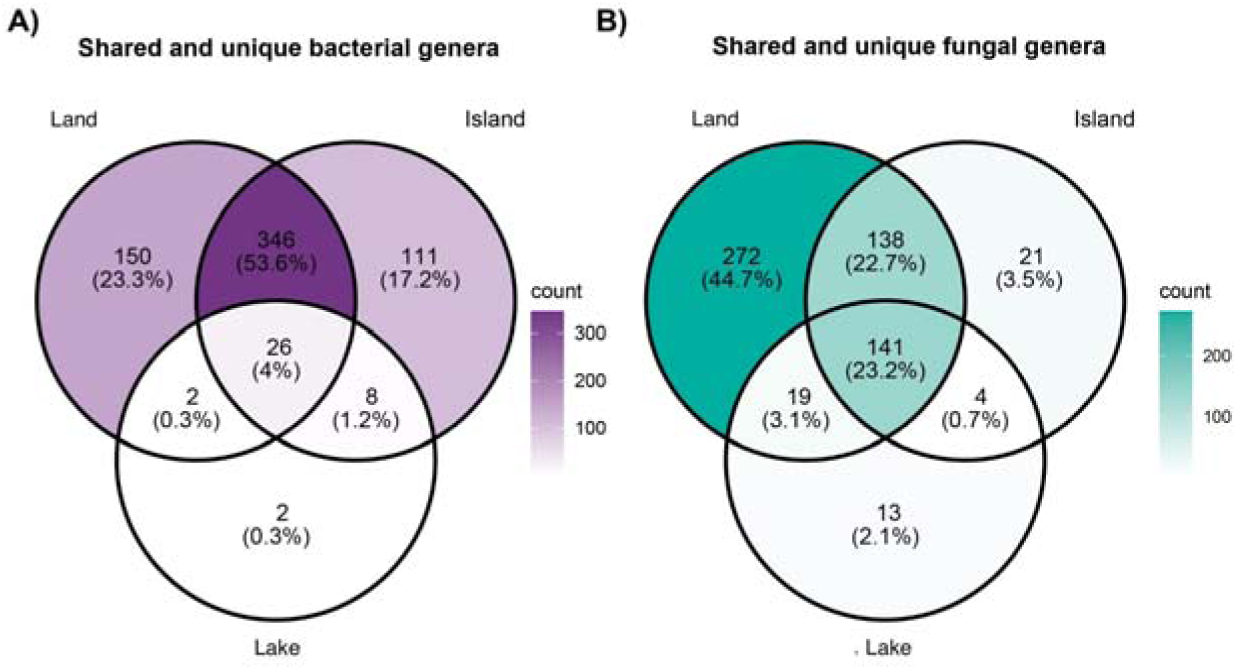
Shared and unique bacterial and fungal genera across habitats. Venn diagrams show the number and percentage of genera unique to or shared among mainland soil (Land), island soil (Island), and lake water (Lake) for (A) bacteria and (B) fungi. Percentages are calculated relative to the total number of bacterial or fungal genera detected, respectively. Color intensity increases with the number of genera detected in each category.

Fungal communities showed considerably greater overlap across habitats. In total, 141 genera (23.2% of all detected fungal genera) occurred in all three habitats and 138 (22.7%) were shared exclusively between Land and Island soil (**Figure 4B**). Nevertheless, the largest fungal fraction consisted of genera unique to Land (272 genera, 44.7%), whereas only 21 genera (3.5%) were unique to Island and 13 (2.1%) to Lake samples. Thus, although fungal genera showed greater overlap across the terrestrial-aquatic boundary than bacterial genera, Island communities predominantly shared genera with terrestrial communities rather than forming a distinct transitional assemblage between Land and Lake.

### Genus overlap changes with distance from the shoreline

To determine whether genus-level overlap also varied spatially within terrestrial and aquatic habitats, we compared communities among Shore, Middle, and Furthest sampling positions (**Figure 5**). This analysis revealed contrasting distance-related patterns between terrestrial and aquatic communities. Within terrestrial habitats, most bacterial genera (189, 36.1%) were detected across all three distances from the shoreline. A further 148 genera (28.2%) were shared exclusively between Shore and Middle sites (**Figure 5A**). The number of bacterial genera unique to individual distance categories decreased with increasing distance from the shoreline, from 106 at Shore to 62 at Middle, with no genera detected exclusively at Furthest sites. In Lake samples, considerably fewer bacterial genera were shared among distance categories: only eight genera (21.1%) occurred at Shore, Middle and Furthest sites, while the largest fraction (23; 60.5%) was detected exclusively at Furthest sites (**Figure 5B**).

**Figure 5.**
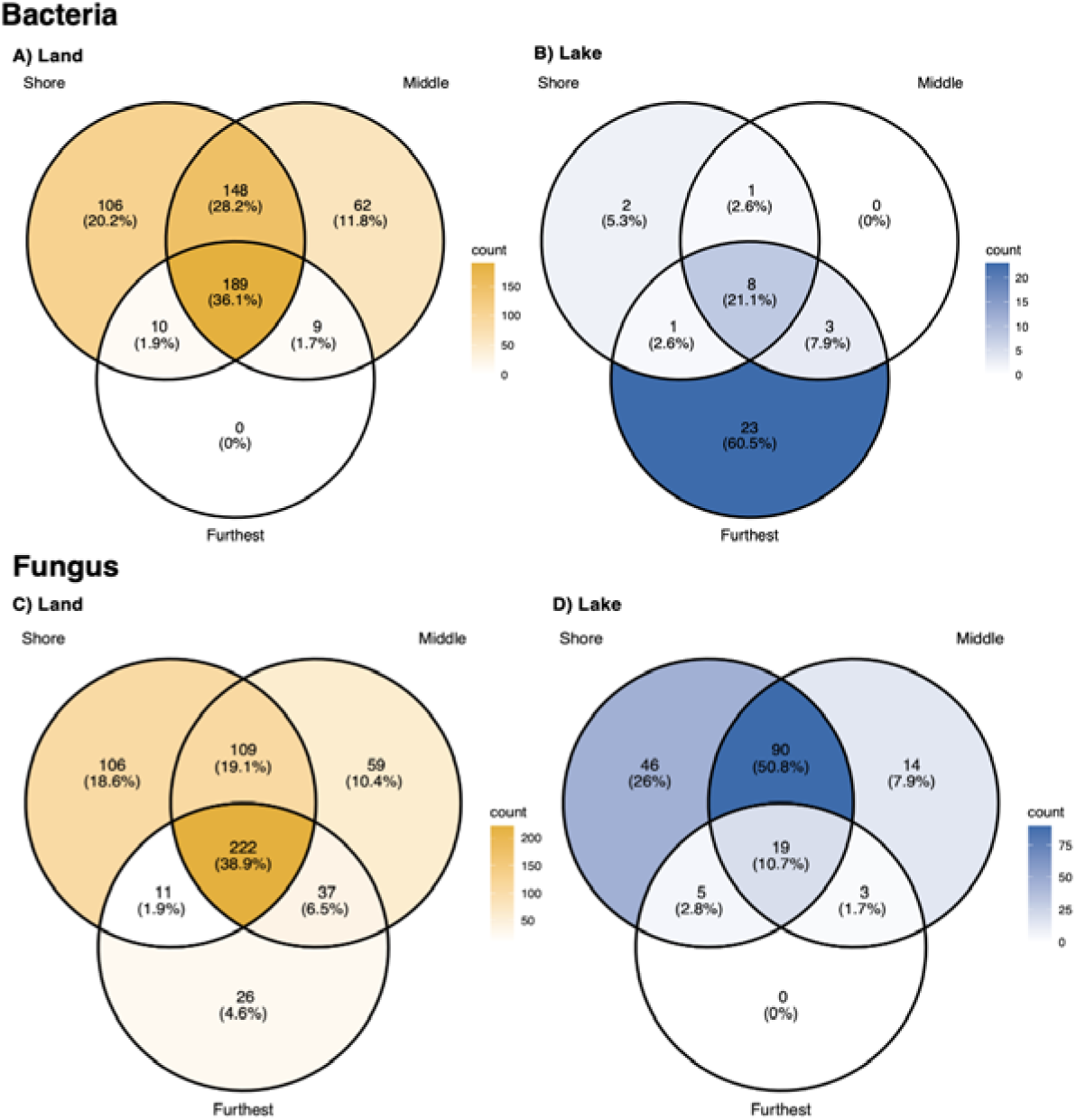
Shared and unique bacterial and fungal genera across distances from the shoreline. Venn diagrams show the number and percentage of genera unique to or shared among Shore, Middle and Furthest sampling positions within A) bacterial Land and B) bacterial Lake, C) fungal Land and D) fungal Lake communities. Percentages are calculated relative to the total number of genera detected within each habitat and taxonomic group. The color intensity increases with the number of genera detected.

Fungal communities in terrestrial habitats showed a broadly similar pattern. Most fungal genera (222; 38.9%) were shared across all three distances from the shoreline, while the number of genera unique to each distance decreased from Shore (106) to Middle (59) and Furthest (26; **Figure 5C**). But unlike bacteria, fungal communities in the Lake showed extensive overlap between Shore and Middle sites, which shared 90 genera (50.8%), while a further 19 genera (10.7%) occurred across all three distances. (**Figure 5D**). No fungal genera were detected exclusively at Furthest sites.

### Lake samples show strong dominance of a small number of abundant genera

A total of 3,969,692 bacterial reads corresponding to 8,011 ASVs were recovered from Land samples, 955,579 reads corresponding to 3,752 ASVs from Island samples, and 6,518,578 reads corresponding to 215 ASVs from Lake samples. Thus, despite the higher sequencing depth in Lake samples, considerably fewer bacterial ASVs were recovered, with sequencing reads concentrated among a small number of abundant genera. Notably, a single ASV assigned to the genus *Pseudomonas* accounted for 1,263,568 reads, corresponding to 19.4% of all bacterial reads recovered from Lake samples. Across all habitats, the 20 most abundant bacterial genera accounted for 82.31% of bacterial reads, with the remaining 17.69% pooled as “Others” (**Figure 6A**).

**Figure 6.**
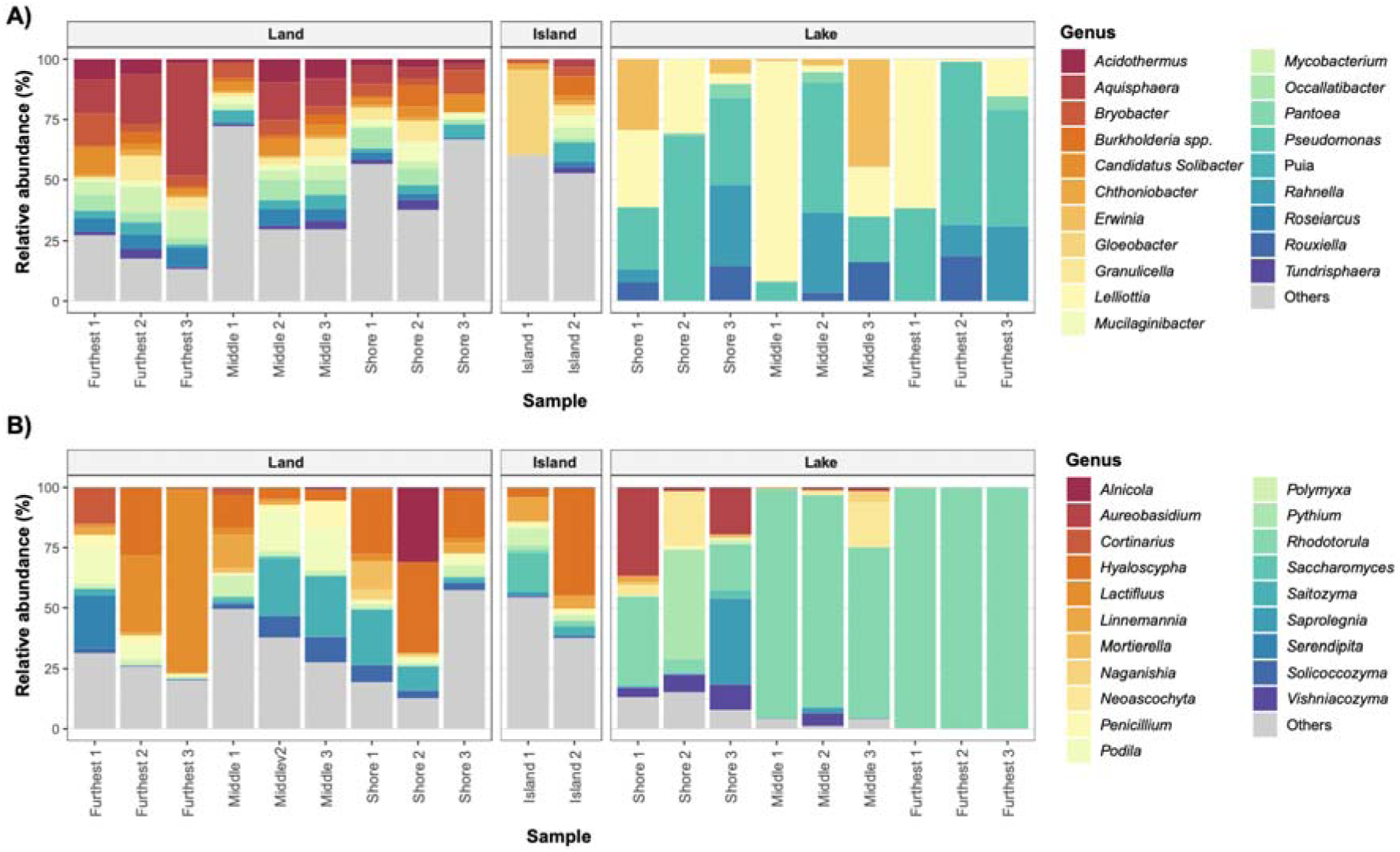
Relative abundance of 20 most abundant A) bacterial and B) fungal genera across samples. Bars represent sample sites (n = 2 replicates) and are grouped by habitat, i.e., Land, Island and Lake. Colours represent the 20 most abundant genera; with remaining genera pooled as ‘Others’. Within Land and Lake groups, samples are ordered by Furthest, Middle and Shore positions.

A similar pattern was observed for fungi. A total of 7,757,822 fungal reads corresponding to 4,254 ASVs were recovered from Land samples, 1,259,268 reads corresponding to 1,336 ASVs from Island samples, and 18,600,858 reads corresponding to 488 ASVs from Lake samples. Despite the substantially greater sequencing depth in Lake samples, reads were again concentrated among relatively few ASVs. A single ASV assigned to the genus *Rhodotorula* accounted for 7,327,125 reads, representing 39.4% of all fungal reads recovered from Lake samples. Across all habitats, the 20 most abundant fungal genera accounted for 87.47% of the fungal reads, with the remaining 12.53% pooled as “Others” (**Figure 6B**).

### Ecological roles differ among habitats

Given the strong habitat-associated differences in taxonomic composition, we next investigated whether these patterns were accompanied by differences in potential ecological roles. Bacterial ecological functions and fungal ecological guilds were inferred from ASV taxonomic assignments using FAPROTAX and FUNGuild, respectively.

For bacteria, inferred functions differed markedly among habitats (Figure 7). Chemoheterotrophy, particularly aerobic chemoheterotrophy, was highly represented across Land, Island and Lake samples. Lake samples were additionally characterized by high relative abundances of anaerobic chemoheterotrophy and fermentation. In contrast, several functions associated with nitrogen, methane, and sulfur cycling were predominantly represented in terrestrial samples. Photoautotrophic functions, including oxygenic photoautotrophy and functions associated with photosynthetic cyanobacteria, showed high relative abundance at one Island site, coinciding with the detection of *Cyanobacteriia*.

**Figure 7.**
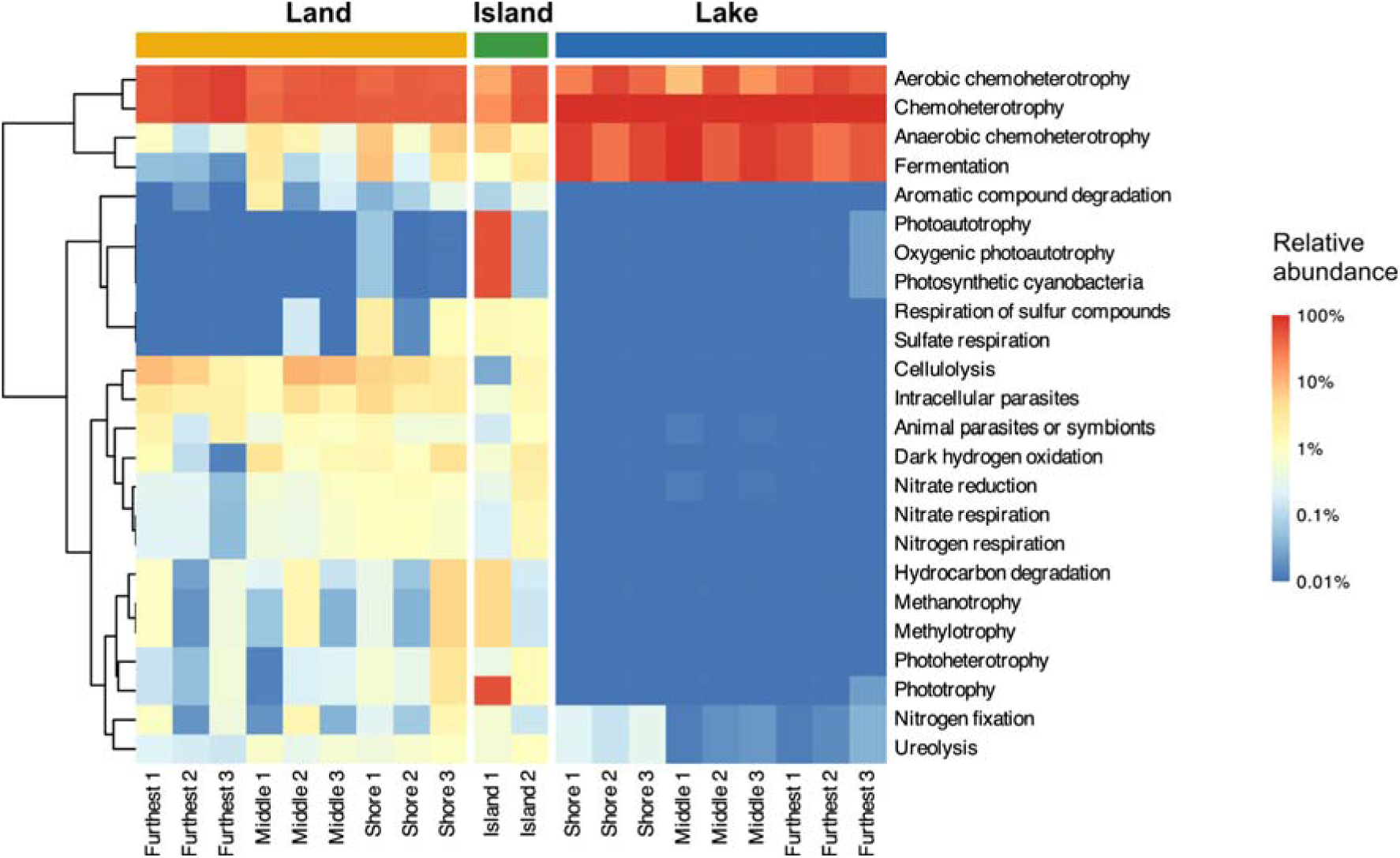
Taxonomically inferred bacterial functions across habitats. Heatmap showing the relative abundance of FAPROTAX functional categories across Land, Island and Lake sampling sites. Rows represent functional categories reaching ≥1% relative abundance in at least one site, and columns represent individual sampling sites, with replicate read counts combined within each site. Within Land and Lake habitats, columns are grouped by distance from the shoreline (Furthest, Middle and Shore), with sampling sites closest to the land-water boundary positioned toward the center of the figure. Colours show relative abundance on a logarithmic scale. Rows are hierarchically clustered according to similarity in their abundance profiles across sites.

Inferred fungal guild composition also showed clear differences among habitats (**Figure 8**). Terrestrial samples (Land and Island) were characterized by relatively high representation of mycorrhizal and saprotrophic guilds, particularly ectomycorrhizal, ericoid mycorrhizal and wood-saprotroph assignments. Island samples showed broadly similar guild profiles to mainland soil, plus high contributions from undefined saprotrophs and plant-associated guilds. Several guilds, including lichen and orchid-mycorrhizal associations, were largely restricted to terrestrial samples. In contrast, Lake samples showed lower relative abundances of most mycorrhizal and wood-saprotroph guilds, especially at the Furthest sites. Instead, Lake samples displayed higher relative abundances of animal-associated and pathogenic guilds, including animal pathogen, animal symbiotroph, endophyte and plant pathogen assignments. This contrast was especially pronounced at Furthest Lake sites, where these guilds accounted for most of the assigned fungal guild abundance. Overall, taxonomically inferred bacterial functions and fungal guilds reflected the previously observed differentiation between terrestrial and aquatic communities. Island communities again exhibited profiles more similar to terrestrial mainland communities than to aquatic communities, indicating that the habitat-associated taxonomic patterns were accompanied by corresponding differences in potential ecological roles.

**Figure 8.**
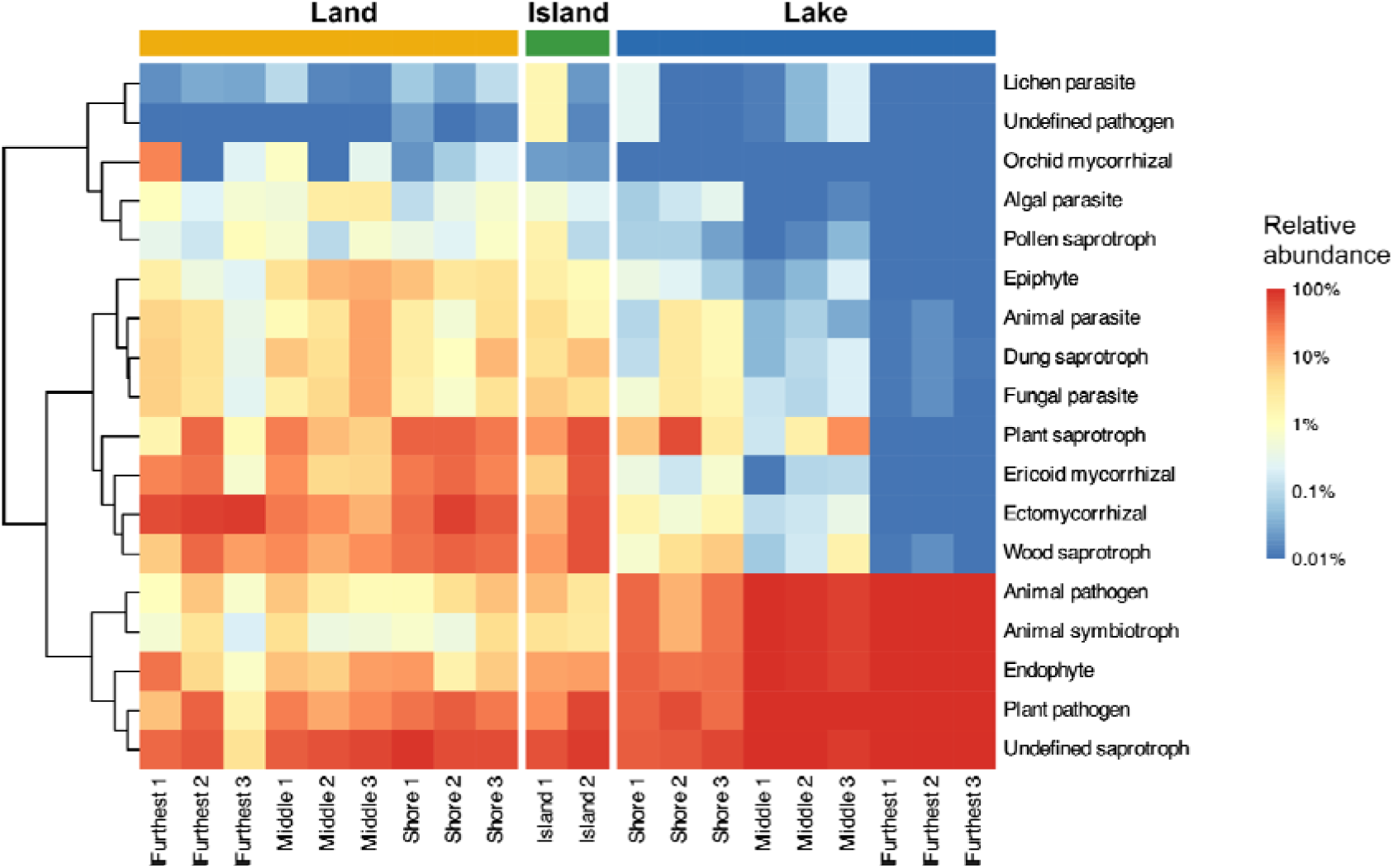
Taxonomically inferred fungal guilds across habitats. Heatmap showing the relative abundance of FUNGuild functional categories across Land, Island and Lake sampling sites. Rows represent functional categories reaching ≥1% relative abundance in at least one site, and columns represent individual sampling sites, with replicate read counts combined within each site. Within Land and Lake habitats, columns are grouped by distance from the shoreline (Furthest, Middle and Shore), with sampling sites closest to the land-water boundary positioned toward the center of the figure. Colours show relative abundance on a logarithmic scale. Rows are hierarchically clustered according to similarity in their abundance profiles across sites.

### Bacterial and fungal genera show strong cross-kingdom co-occurrence patterns

Having identified habitat-associated differences in taxonomic composition and inferred ecological roles, we next examined associations among consistently detected bacterial and fungal genera using a co-occurrence network. To reduce associations driven by rare taxa or individual samples, only genera reaching a relative abundance >1% in at least two sampling sites were retained. The resulting network contained 38 significant associations, both positive and negative, within and between bacterial and fungal communities (**Figure 9**).

**Figure 9.**
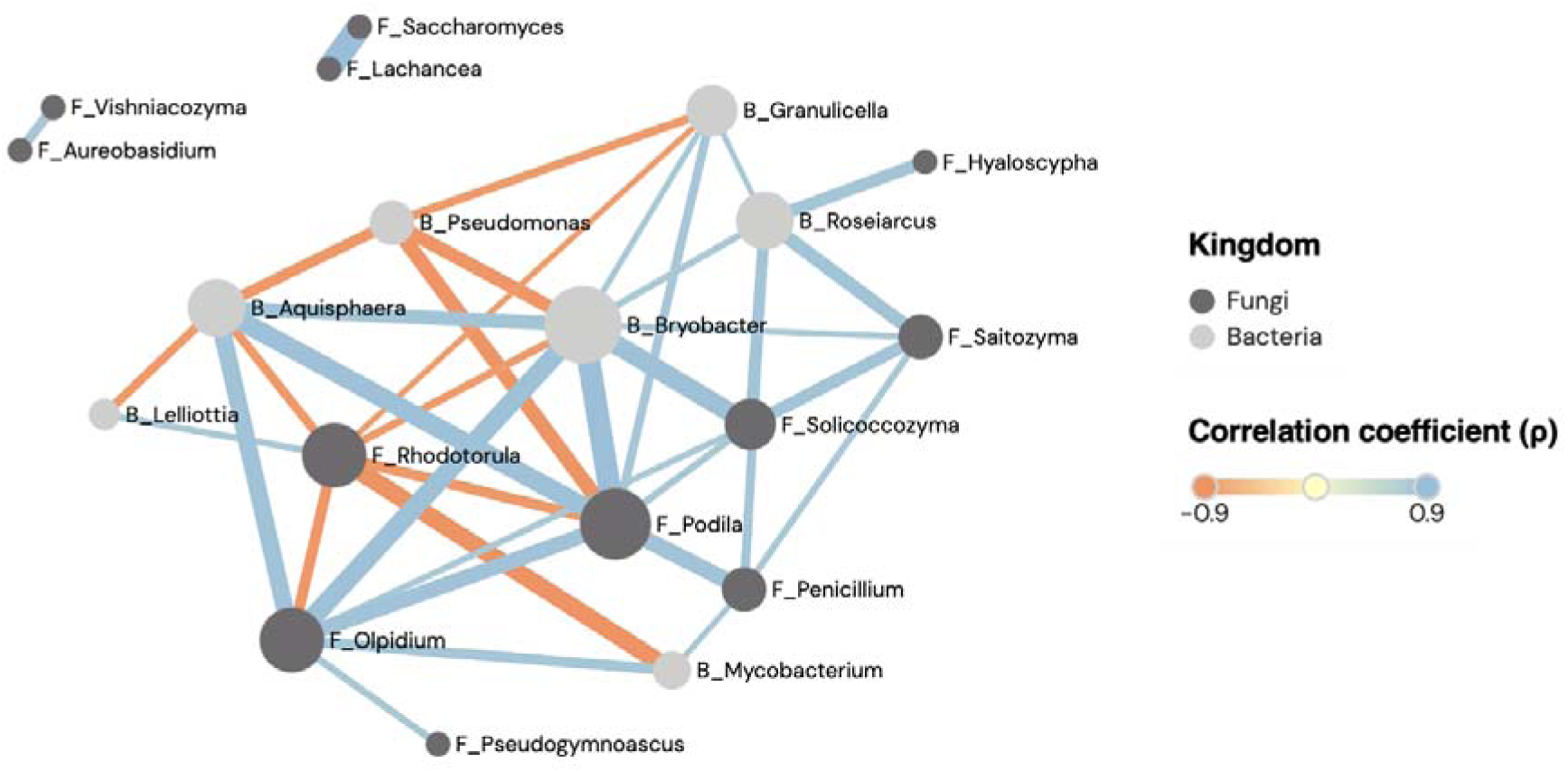
Bacterial-fungal co-occurrence network across samples. The network shows significant Spearman correlations (|ρ| ≥ 0.8 and p_adj_ < 0.05, with Benjamini-Hochberg correction) among bacterial and fungal genera found across sampling sites. Only genera reaching >1% relative abundance in at least two sites were included. Nodes represent genera, with node colour indicating taxonomic group (bacteria or fungi). Prefixes B_ and F_ denote bacterial and fungal genera, respectively. Node size is proportional to connectivity. Edges represent significant positive or negative associations between genera, with edge color indicating the direction and the magnitude of Spearman’s ρ. Edge width is proportional to ρ.

Several genera occupied highly connected positions in the network, including the bacterial genera *Bryobacter* and *Aquisphaera* (**Figure 9**, light grey) and the fungal genera *Podila*, *Olpidium* and *Rhodotorula* (**Figure 9**, dark grey). The strongest negative cross-kingdom association was observed between *Mycobacterium* and *Rhodotorula* (Spearman’s ρ = –0.89, **Suppl. Table S5**), whereas *Bryobacter* and *Podila* showed a strong positive association (ρ = 0.93). Within bacteria, *Bryobacter* and *Pseudomonas* showed the strongest negative association (ρ = –0.86), while *Aquisphaera* and *Bryobacter* were strongly positively associated (ρ = 0.87). Among fungi, *Olpidium* and *Rhodotorula* showed a strong negative association (ρ = –0.84), whereas *Lachancea* and *Saccharomyces* showed the strongest positive association (ρ = 0.95).

Together, these results reveal strong patterns of positive and negative co-occurrence both within and between bacterial and fungal communities These associations may reflect shared or contrasting habitat preferences as well as potential ecological interactions among taxa.

Given the strong bacterial-fungal associations and the influences of the habitat in microbial diversity and inferred ecological roles, we examined whether co-occurrence patterns also varied among habitats (**Suppl. Table S5**). We calculated correlation coefficients (ρ) separately for Lake, Island and Land samples. Of the 38 significant associations detected across the full dataset, 12 involved pairs of genera that were both not detected in Lake samples, indicating that these network associations were restricted to terrestrial samples. These included several associations among bacterial genera, such as *Aquisphaera-Bryobacter* and *Bryobacter-Roseiarcus*, as well as associations involving fungal genera such as *Podila*. Furthermore, the fungal-fungal association between *Olpidium* and *Pseudogymnoascus*, for which both genera were not detected in Lake samples, was also the only association for which both genera were undetected in Island samples. All identified associations involved genera detected in Land.

In contrast, some associations showed strong positive correlations in all three habitats. For example, *Lachancea-Saccharomyces* showed strong positive associations in both Land (ρ = 0.967), Lake (ρ = 0.883) and Island (ρ = 1). Other associations differed markedly in correlation strength between habitats, despite both genera being detected in all habitats, such as *Aureobasidium-Vishniacozyma* (ρ = 0.271 in Land vs. ρ = 0.851 in Lake and ρ = 1 in Island).

Together, these results revealed both consistent and habitat-dependent patterns of co-occurrence. Some genus pairs remained strongly positively correlated across Land, Island and Lake samples, whereas others showed marked differences in correlation strength among habitats, even when both genera were detected. Thus, habitat-associated variation extended beyond community composition and inferred ecological roles to the patterns of association among microbial genera.

## Discussion

Strong habitat-associated differentiation of bacterial and fungal communities emerged across a land-water interface of only 350 m. Despite the close geographic proximity, Land, Island, and Lake habitats supported distinct microbial assemblages, highlighting the ecological heterogeneity that can occur over short distances. Previous studies have shown that microbial communities can differ markedly between neighbouring habitats, including across terrestrial-freshwater transitions (Martiny *et al*., 2006; O’Malley, 2008; Monard *et al*., 2016). Our findings extend these observations to a particularly small spatial scale, where community composition was strongly associated with habitat identity. In contrast, neither bacterial nor fungal community composition varied significantly along the southwest-northeast sampling axis parallel to the shoreline. This suggests that geographic distance alone was not a major determinant of the observed patterns. Instead, the pronounced differences we found between and within terrestrial and aquatic samples are consistent with the hypothesis that local habitat conditions contribute to maintaining distinct microbial assemblages despite their close geographic proximity. Although our study did not directly quantify dispersal or environmental selection, these findings highlight the potential importance of habitat-associated processes in structuring microbial communities across fine-scale environmental boundaries.

The microbial communities on the two small islands we sampled provide an interesting perspective on the relationship between habitat identity and geographic connectivity. Despite being surrounded by lake water, Island communities were more similar to those found in mainland soil than to those of the surrounding aquatic habitat. Bacterial and fungal community composition did not differ significantly between Land and Island samples, whereas both were significantly differentiated from Lake samples. This pattern was also reflected in genus-level analyses, which revealed a substantially greater taxonomic overlap between Island and Land than between island soil and lake water. Rather than supporting transitional communities between terrestrial and aquatic environments, the islands therefore appeared to maintain predominantly terrestrial microbial assemblages. Interestingly, bacterial genus richness was even higher on the islands than on the mainland, suggesting that similarity in overall community composition does not necessarily imply identical levels of local diversity. Similar associations between island and mainland microbial communities have been reported in studies of microbial island biogeography (Li *et al*., 2020). Together, these findings suggest that local habitat characteristics may contribute to maintaining terrestrial-like communities even on small islands embedded within an aquatic environment.

In this aquatic environment, the bacterial and fungal diversity was lower (**Figure 2**, **3**) consistently with previous studies reporting higher microbial richness in soils than in freshwater environments. This potentially reflects the greater heterogeneity of microhabitats and resources available in terrestrial environments (Sieber *et al*., 2020; Saona *et al*., 2024). The strong representation of *Pseudomonas* and *Rhodotorula* in Lake samples (**Figure 6A**, **6B**) is also consistent with the occurrence of these genera in freshwater environments. *Pseudomonas* can exploit a broad range of organic substrates (Flynn *et al*., 2017) and has previously been reported as abundant in aquatic habitats (Ponce-Soto *et al*., 2015). Similarly, *Rhodotorula* occurs in freshwater environments, including oligotrophic lakes, where its ability to utilize a broad range of carbon sources and tolerate environmental stress may contribute to its persistence (Brandão *et al*., 2011). Some *Rhodotorula* species also produce extracellular enzymes involved in the degradation of organic compounds, suggesting a potential role in aquatic carbon cycling (Lario *et al*., 2015).

Alongside these ecological explanations, the YNB-raffinose enrichment applied to Lake samples may also have contributed to the observed patterns. By favouring microorganisms capable of proliferating under the provided conditions, enrichment could have increased the relative representation of certain genera while reducing the abundance of others, thereby contributing to the lower observed richness observed in aquatic samples (Eilers, Pernthaler and Amann, 2000; Sampaio and Gonçalves, 2008; Pinto *et al*., 2025). Land and Island samples, in contrast, showed high genus richness and were processed using the same procedures, allowing their comparison without the additional influence of the enrichment applied to Lake samples. The observed patterns may therefore reflect a combination of ecological differences and enrichment effects, although their relative contributions cannot be distinguished with the present data.

Beyond the differences among habitats, bacterial and fungal communities showed contrasting patterns of spatial differentiation with distance from the shoreline. Bacterial community composition did not vary significantly among Shore, Middle and Furthest positions within either terrestrial or aquatic habitats. In contrast, fungal community composition varied significantly with shoreline position in both habitats, and the significant interaction between habitat and shoreline position indicated that these spatial patterns differed between Land and Lake. Although individual pairwise comparisons among positions were not significant after correction for multiple testing, these results suggest that fungal community composition may be more strongly structured along local shoreline gradients than bacterial community composition. On Land, spatial variation coincided with changes in vegetation from predominantly deciduous forest at Shore and Middle sites to coniferous forest at Furthest sites (**Figure 1**). Vegetation composition can influence fungal communities through differences in litter inputs, plant associations and soil properties, while associated variation in moisture and other abiotic conditions may further contribute to community differentiation (Bahnmann *et al*., 2018; Philippot *et al*., 2024; Lyu *et al*., 2026). Mycorrhizal fungi provide a relevant example, as their associations with plant roots may link community composition to changes in the availability and identity of host plants (Saona *et al*., 2024). Their greater representation in terrestrial samples and reduced representation in Lake samples (**Figure 8**) are also consistent with the contrasting availability of forest-associated hosts across these habitats. These environmental characteristics were not quantified directly, but provide plausible explanations for the observed spatial patterns.

Within Lake samples, fungal communities also differed significantly with shoreline position, with a relatively large compositional difference between Shore and Furthest sites. Although fungal alpha diversity tended to be lower at Furthest sites, pairwise differences were not statistically significant (**Figure 2**, **3**). One possible explanation is that aquatic fungal communities closer to shore receive greater inputs of terrestrial organic matter, plant-associated microorganisms and fungal spores through litterfall or runoff (Strayer and Findlay, 2010; Hayer *et al*., 2022). Such inputs could contribute to spatial variation in the genera detected within the aquatic habitat. Consistent with this possibility, most fungal genera detected in Lake samples occurred at Shore or Middle sites, while none were detected exclusively at Furthest sites (**Figure 5D**). However, shoreline position also coincided with differences in water depth, and other unmeasured environmental factors could contribute to the observed patterns. The underlying mechanisms therefore remain unresolved.

Interestingly, fungal communities showed substantially greater genus-level overlap between terrestrial and aquatic habitats than bacterial communities (**Figure 4**). Nevertheless, this greater overlap did not translate into similar overall community composition, as Land and Lake fungal communities remained significantly differentiated. This suggests that shared genera may differ in their relative representation across habitats, alongside differences in the occurrence of habitat-associated genera. Soils can act as reservoirs of microorganisms that enter adjacent aquatic environments (Crump, Amaral-Zettler and Kling, 2012; Ruiz-González, Niño-García and Del Giorgio, 2015). However, the detection of a genus in both terrestrial and aquatic samples does not necessarily demonstrate successful colonization or establishment. Amplicon sequencing detects DNA rather than active growth, and some shared taxa may represent transient material transported between habitats. Furthermore, the enrichment procedure may have influenced which aquatic taxa were subsequently detected. Thus, differences in taxonomic overlap between bacteria and fungi may reflect a combination of ecological characteristics, transport processes and methodological effects, rather than differences in dispersal alone.

The pronounced differentiation among habitats was also reflected in the ecological roles inferred from their constituent taxa, adding an ecological dimension to the observed taxonomic patterns. Terrestrial bacterial communities showed higher representation of several functional categories associated with nitrogen, methane and sulfur cycling, whereas Lake samples showed higher relative abundances of anaerobic chemoheterotrophy and fermentation (**Figure 7**). Similar differences in predicted functional profiles between soil and aquatic communities have previously been reported (Zhang *et al*., 2024). For fungi, terrestrial communities were characterized by greater representation of mycorrhizal and wood-saprotrophic guilds, consistent with the presence of forest vegetation and associated organic substrates. In contrast, Lake samples showed higher relative abundances of several animal-associated, endophytic, and pathogenic guilds (**Figure 8**). Comparable differences in fungal guild composition have also been observed between terrestrial and aquatic environments (Mircea *et al*., 2024). Island communities again showed profiles broadly resembling those of mainland soil. These patterns suggest that habitat-associated differences in taxonomic composition are accompanied by differences in predicted ecological roles. However, because these functional annotations were inferred from taxonomic composition rather than measured directly, they should be interpreted as potential ecological capacities rather than evidence of functional activity.

Finally, the co-occurrence network identified several strong positive and negative associations among bacterial and fungal genera, both within and between kingdoms (**Figure 9**). Some associations involved genera detected exclusively in terrestrial samples, whereas others involved genera occurring across both terrestrial and aquatic habitats. For example, *Lachancea* and *Saccharomyces* showed the strongest positive correlation across the full dataset, while *Mycobacterium* and *Rhodotorula* showed the strongest negative correlation; *Mycobacterium* being exclusively detected in the soil, whereas *Rhodotorula* was more abundant in the water. These associations among microbial genera may reflect shared or contrasting habitat preferences, but could also arise from ecological interactions among taxa. Functional annotations of taxa involved in the co-occurrence network revealed associations among taxa with both overlapping and distinct annotated ecological roles (**Suppl. Table S5**). Most bacterial-bacterial associations involved genera annotated as aerobic chemoheterotrophs, such as *Aquisphaera*, *Bryobacter*, and *Pseudomonas*. Bacteria-fungal associations frequently involved bacteria annotated as chemoheterotrophs and fungi assigned to saprotrophic or plant-associated guilds. As for fungal-fungal associations, fungal taxa involved were mostly assigned to saprotrophic guilds, including *Saccharomyces*, *Saitozyma*, *Solicoccozyma*, and *Vishniacozyma*, although several genera received multiple or broad FUNGuild assignments. However, correlations based on relative abundances cannot establish direct biological interactions, particularly when communities differ strongly in their composition across habitats. The observed associations should therefore be interpreted as exploratory patterns that may help identify candidate microbial relationships for future investigation.

Overall, our findings reveal pronounced habitat-associated differentiation in bacterial and fungal communities across a geographically restricted land-water interface. The predominantly terrestrial character of island communities, despite their location within the lake, suggests that geographic proximity alone does not necessarily lead to microbial community homogenization. At the same time, the contrasting spatial patterns and taxonomic overlap of bacteria and fungi indicate that microbial groups may respond differently to local environmental gradients and habitat boundaries. Although differences in sample processing limit conclusions about the original aquatic communities, the results highlight the importance of considering habitat identity, spatial context and organismal differences when investigating microbial community structure at fine spatial scales. Future studies combining direct environmental sequencing, measurements of local abiotic conditions and estimates of microbial dispersal would help determine the mechanisms maintaining these community boundaries.

## Acknowledgements

We thank Pablo Villarreal for his comments on the manuscript. The work was funded by the Swedish Research Council (grant 2022-03427) and the Knut and Alice Wallenberg Foundation (2017.0163) to RS. The field work was made possible by a grant from the Albert & Maria Bergströms Foundation to JP and CH. The authors acknowledge support from the National Genomics Infrastructure in Stockholm funded by the Science for Life Laboratory, and the SNIC/Uppsala Multidisciplinary Center for Advanced Computational Science for assistance with sequencing and access to the UPPMAX computational infrastructure. Computation and data handling were enabled by resources in project NAISS 2023/22-116 provided by the National Academic Infrastructure for Supercomputing in Sweden (NAISS) at UPPMAX, funded by the Swedish Research Council through grant agreement no. 2022-06725.

