## Supplementary material for "Strong habitat-associated differentiation of microbial communities across a fine-scale land-water interface": Supp. Fig S1 and Table S1

**Supp. Table S1.** Primer sequences for PCR1

| Name | Forward sequence (5′-3′) | Reverse sequence (5′-3′) |
| --- | --- | --- |
| NGI overhangs - ITS1 | ACACTCTTTCCCTACACGACGCTCTTCCGATCT - TCCGTAGGTGAACCTGCGG | GTGACTGGAGTTCAGACGTGTGCTCTTCCGATCT - GCTGCGTTCTTCATCGATGC |
| NGI overhangs - 16S V3-V4 | ACACTCTTTCCCTACACGACGCTCTTCCGATCT - CCTACGGGNGGCWGCAG | GTGACTGGAGTTCAGACGTGTGCTCTTCCGATCT - GACTACHVGGGTATCTAATCC |

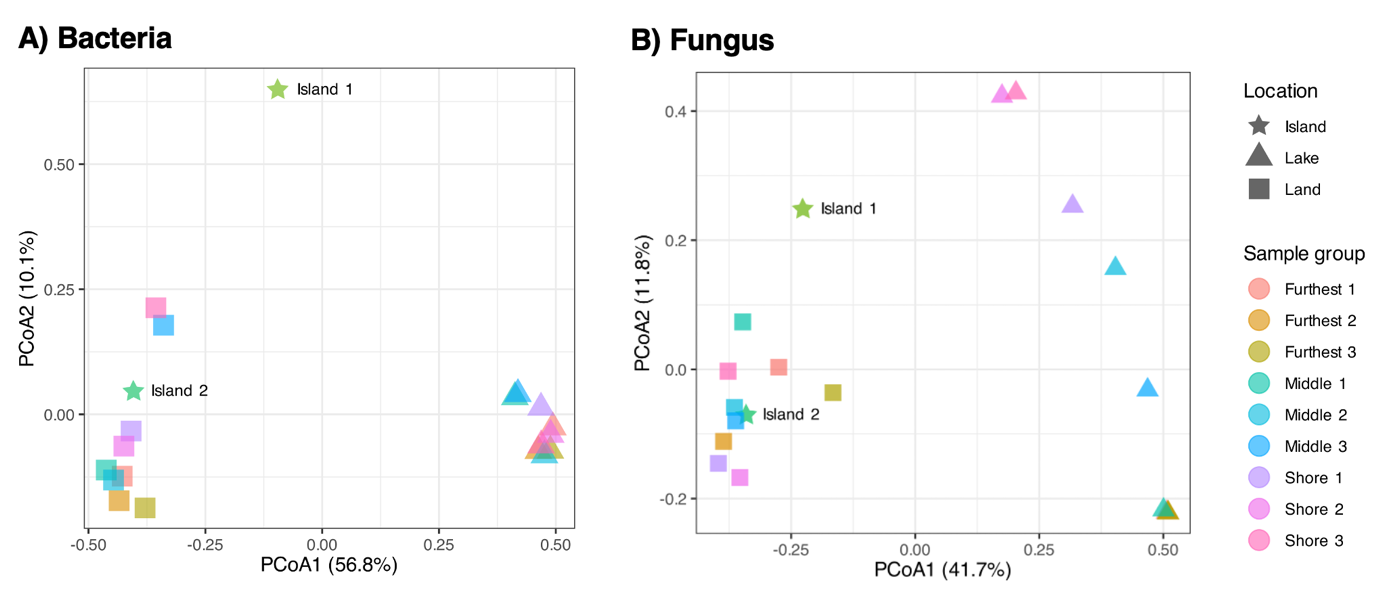

**Supp. Figure S1.** Principal coordinates analysis (PCoA) of A) Bacterial and B) Fungal gene community composition based on Bray-Curtis dissimilarities. Points represent individual samples, with colours indicating sampling site and shapes indicating sample habitats (Land, Island, and Lake).
